# Development of a Recombinant Single-Cycle Influenza Viral Vector as an Intranasal Vaccine against SARS-CoV-2 and SARS-like Betacoronaviruses

**DOI:** 10.1101/2025.11.28.691177

**Authors:** Jonathan Munro, Diana Melnyk, Madeeha Afzal, Lisa Schimanski, Alexander A. Cohen, Jennifer R. Keeffe, Pamela J. Bjorkman, William S. James, Alain R. Townsend, Tiong Kit Tan

## Abstract

The COVID-19 pandemic has demonstrated the detrimental potential of zoonotic coronavirus transmission to human populations. Effective vaccines capable of eliciting immunity to SARS-CoV-2 have been pivotal in mitigating the spread of the virus. In this study, we describe the generation of a non-replicating pseudotyped influenza A virus (S-FLU), where the native haemagglutinin (HA) sequence is replaced with the coding sequence of either a membrane-anchored form (TM) or secretory form (Sec) of the receptor-binding domain (RBD) of the ancestral SARS-CoV-2 Wuhan (S-RBD Wuhan). We showed that both S-RBD-TM and S-RBD-Sec viruses can be generated via reverse genetics and grown to high titre. Intranasal immunisation in mice with S-RBD-TM elicits robust serum binding and neutralisation activity against SARS-CoV-2, superior to S-RBD-Sec. Furthermore, we demonstrate that a heterologous prime-boost immunisation regime in mice with S-RBD-TM Wuhan and S-RBD-TM BM48-31 (a distant Clade 3 SARS-like betacoronavirus (sarbecovirus)) increases antibody breadth against mismatched sarbecoviruses compared to homologous prime-boost with S-RBD-TM Wuhan. These results suggest that S-RBD-TM is a promising intranasal vaccine candidate against SARS-CoV-2 and may offer potential as a broadly protective sarbecovirus vaccine.

## Introduction

The global pandemic of 2020 was triggered by the emergence of SARS-CoV-2, a zoonotic SARS-like betacoronavirus (sarbecovirus) responsible for the respiratory disease COVID-19 [1, 2]. SARS-CoV-2 caused over 6 million reported deaths, with estimates of the true toll reaching 17.2 million [3]. Whilst vaccines have proven successful in reducing morbidity and mortality, several variants of concern (VOCs) have emerged, exhibiting increased transmissibility, disease severity, and immune evasion [4–8], posing challenges to vaccine efficacy [9–12].

Prior to SARS-CoV-2, six members of the *Coronaviridae* family spilled over to humans from animal reservoirs [13–17]. While four are endemic and cause non-severe symptoms [13], SARS-CoV-1 and MERS-CoV caused high mortality and morbidity [18–20]. Instances of simultaneous infection with SARS-CoV-2 and MERS-CoV in humans in the Middle East have been reported [21], with both viruses infecting type-II alveolar cells and utilising identical transcription regulatory sequences [1, 22]. Furthermore, the identification of numerous coronaviruses in bats, civets, and pangolin that share similar genomes with SARS-CoV-2 and utilise the human ACE2 as a host cell receptor highlights their potential to spill over into humans [2, 15, 23]. Therefore, there is great concern and apprehension regarding the potential for future zoonotic coronavirus spillover events that could precipitate new pandemics [24–27]. This highlights the importance of being proactive rather than reactive, by developing broader spectrum sarbecovirus vaccines to control the spread of SARS-CoV-2 variants and to prevent future pandemics from novel zoonotic sarbecoviruses [28, 29].

The sarbecovirus spike (S) glycoprotein consists of S1 and S2 subunits [30]. The S1 subunit is responsible for initial contact with host receptors [31] and contains the receptor-binding domain (RBD), which is highly immunogenic and the site recognised by most neutralising antibodies [32–35]. Monoclonal antibodies targeting the Class 1 and 2 RBD epitopes are generally strain-specific, as these epitopes exhibit considerable sequence variability across sarbecoviruses and SARS-CoV-2 variants. In contrast, cross-neutralising monoclonal antibodies against diverse sarbecoviruses and SARS-CoV-2 VOCs have been isolated that primarily recognise the more conserved 4, 1/4, and portions of the Class 3 and Class 5 epitopes on the RBD [36–43]. These findings indicate that RBD-based vaccines designed to focus immune responses on these conserved regions could provide broad-spectrum protection against sarbecovirus.

We describe in this study the use of a modified single-cycle influenza virus vaccine (S-FLU), shown previously to have produced broad protection against influenza [44, 45], as a potential vaccine against SARS-CoV-2 and other zoonotic sarbecoviruses. Although similar approaches have been previously reported [46–49], S-FLU offers a key advantage as it can be pseudotyped safely with avian haemagglutinin subtypes (e.g., H5 or H7) for which there is minimal pre-existing immunity in the human population, thereby reducing the risk of immune interference with vaccine efficacy. Inactivation of the native haemagglutinin (HA) signal sequence means that S-FLU can only replicate in cell lines transfected to express HA that provide the surface protein for budding viral particles [45]. Briefly, we replaced the native HA coding sequence within the HA viral RNA (vRNA) with either a membrane-anchored form of the SARS-CoV-2 Spike RBD (S-RBD-TM) or a secreted form (S-RBD-Sec), based on the prototypic Wuhan SARS-CoV-2 Spike RBD. We showed that both S-RBD-Sec and S-RBD-TM grew to high titre and induced the expression of RBD *in vitro* upon infection. We showed that S-RBD-TM, when given intranasally, elicited robust serum antibody response against SARS-CoV-2. We then tested if a heterologous prime-boost regime with a genetically distant sarbecovirus RBD can improve the breadth of immune response. We produced S-RBD-TM BM48-31 (a Clade 3 sarbecovirus RBD) and showed that a S-RBD-TM Wuhan/S-RBD-TM BM48-31 prime-boost regime moderately increases antibody breadth against mismatched sarbecoviruses compared to homologous prime-boost with S-RBD-TM Wuhan. Together, our results highlight S-RBD-TM as a promising vaccine candidate for broader protection against sarbecoviruses.

## Results

### Generation and characterisation of X31 H3 S-RBD

To generate S-FLU capable of expressing the SARS-CoV-2 RBD, the coding region of the haemagglutinin in the HA vRNA was replaced with the sequence encoding the RBD from the SARS-CoV-2 Spike based on the original Wuhan strain (HA-RBD). In addition to the HA-RBD segment, these viruses contain the other seven vRNAs from H1N1 A/Puerto Rico/1/1934 (PR8). Two versions of S-RBD were generated; S-RBD-Sec (expressing soluble RBD) and S-RBD-TM (expressing a membrane-anchored form of the RBD). To generate S-RBD-TM, we fused a signal peptide (from H7 Influenza HA) to the N-terminus of the RBD coding sequence, and added a transmembrane domain (TM) and cytoplasmic domain at the C-terminus (sourced from various proteins). The 3’ and 5’ packaging sequences of the HA vRNA were retained to facilitate viral packaging (Figure 1A). The S-RBD-Sec contains the same sequence as the S-RBD-TM but lacks the transmembrane domain to allow secretion. Both S-RBD-Sec and S-RBD-TM were generated using reverse genetics in HEK293T cells. These S-RBD viruses lack a functioning HA vRNA, therefore HA was supplied in *trans* in producer MDCK cells to allow viral packaging. For the S-RBD-TM, we first tested the native PR8 H1 haemagglutinin transmembrane and cytoplasmic domain but failed to rescue any virus (data not shown). We speculated that the PR8 H1 TM and cytoplasmic domain interfered with virion packaging. We then tested alternative TMs: the mouse CD80 TM as previously described [47], the influenza N7 neuraminidase TM (a type II glycoprotein), and the SARS-CoV-2 Spike TM (with the last 18 amino acids at the N-terminus deleted to increase surface expression) [50]. All three constructs permitted successful virus rescue, evident by expression of influenza viral protein and SARS-CoV-2 RBD in infected cells (Supplementary Figure 1). Both SARS-CoV-2 spike and mouse CD80 TM expressed comparable levels of RBD whereas the N7 TM resulted in lower RBD expression. Based on these results, we proceeded with the SARS-CoV-2 Spike TM domain for S-RBD-TM. Rescued S-RBD-TM and S-RBD-Sec seed viruses were then propagated in MDCK-SIAT1 cells stably transfected to express the X31 H3N2 (A/H3N2/Aichi/1/1968) HA on the cell surface (MDCK-X31). The viruses were then harvested, and the infectious titre (CID_50_) was determined as previously described [51, 52].

**Figure 1:**
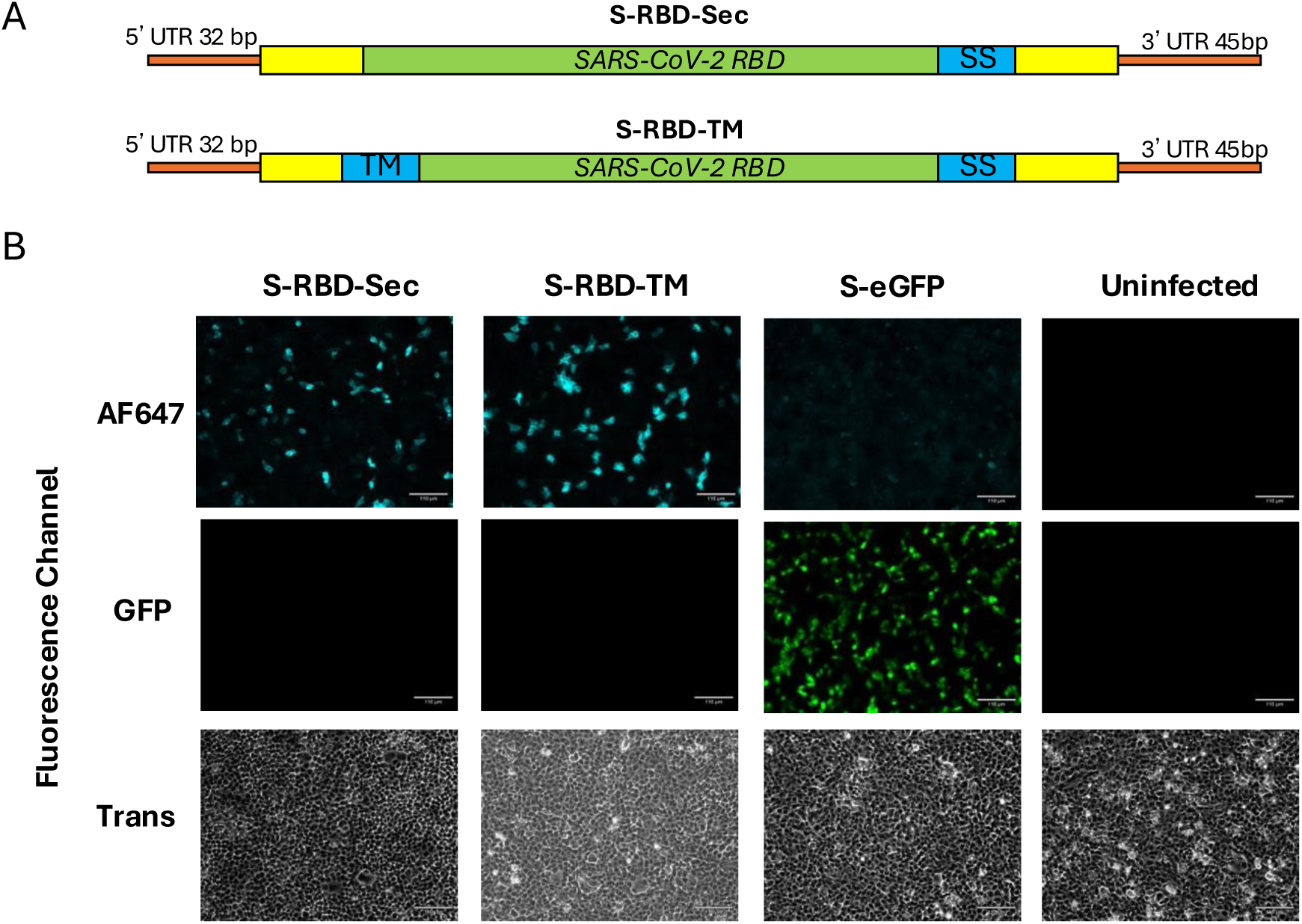
Design and characterisation of S-RBD-Sec and S-RBD-TM. A. Schematic diagram of modified haemagglutinin vRNA to encoding the the membrane-anchored version of RBD or secretory RBD. B. Validation of S-RBD-Sec and S-RBD-TM in cell culture. MDCK cells seeded in 6-well plate were infected with 1e4 CID50 of S-RBD-SEC, S-RBD-TM, S-eGFP (S-Flu expressing GFP) control or remained uninfected. 16 hours post-infection, cells were stained with an RBD-specific IgG1 (EY-6A) followed by a goat anti-human IgG1 labelled with AF647. Cells infected with S-RBD-Sec was permeabilised with 0.5% Triton-X 100 prior to staining. Stained cells were visualised using a fluorescence microscope. Scale bar: 110uM. (UTR: untranslated region, yellow boxes: the 3’ and 5’ packaging sequences of the HA vRNA, SS: signal sequence, TM: transmembrane domain, AF647: Alexa Fluor 647)

To assess RBD expression, MDCK-SIAT1 cells were infected with multiplicity of infection (MOI) 0.01 of the S-RBD-TM and S-RBD-Sec, S-eGFP (an S-FLU expressing GFP), or remained uninfected. After overnight incubation, the infected cells were stained with EY-6A, a class 4 anti-RBD monoclonal antibody known to bind to all sarbecovirus RBDs [33]. The results show that both S-RBD-TM and S-RBD-Sec led to expression of RBD in the infected cells (Figure 1B), confirming successful incorporation and expression of the SARS-CoV-2 RBD.

### Mice vaccinated with S-RBD generate robust systemic anti-RBD antibody responses

To determine if S-RBD-TM and S-RBD-Sec induce an anti-RBD antibody response *in vivo*, mice (female C57BL/6, 5 weeks old) were immunised twice, two weeks apart, intranasally (IN) or intramuscularly (IM) with 5.5e5 CID_50_ of either S-RBD-TM, S-RBD-Sec, or S-eGFP as a mock control (Figure 2A). Sera were collected 2 weeks post-prime and at 3 weeks post-boost. Serum anti-RBD IgG antibody responses were measured using RBD ELISAs (Figure 2B, C) and authentic virus neutralisation assays (Figure D) [53–55]. The data show that the intranasal route induces a generally higher serum anti-RBD binding response than the intramuscular route, and the S-RBD-TM is more immunogenic than the S-RBD-Sec when the same dose is given. In addition, the S-RBD-TM administered intranasally successfully produced neutralising titres against authentic SARS-CoV-2, while the other administrations produced titres below detection. Based on these results, we selected the S-RBD-TM and intranasal route for subsequent immunisations.

**Figure 2:**
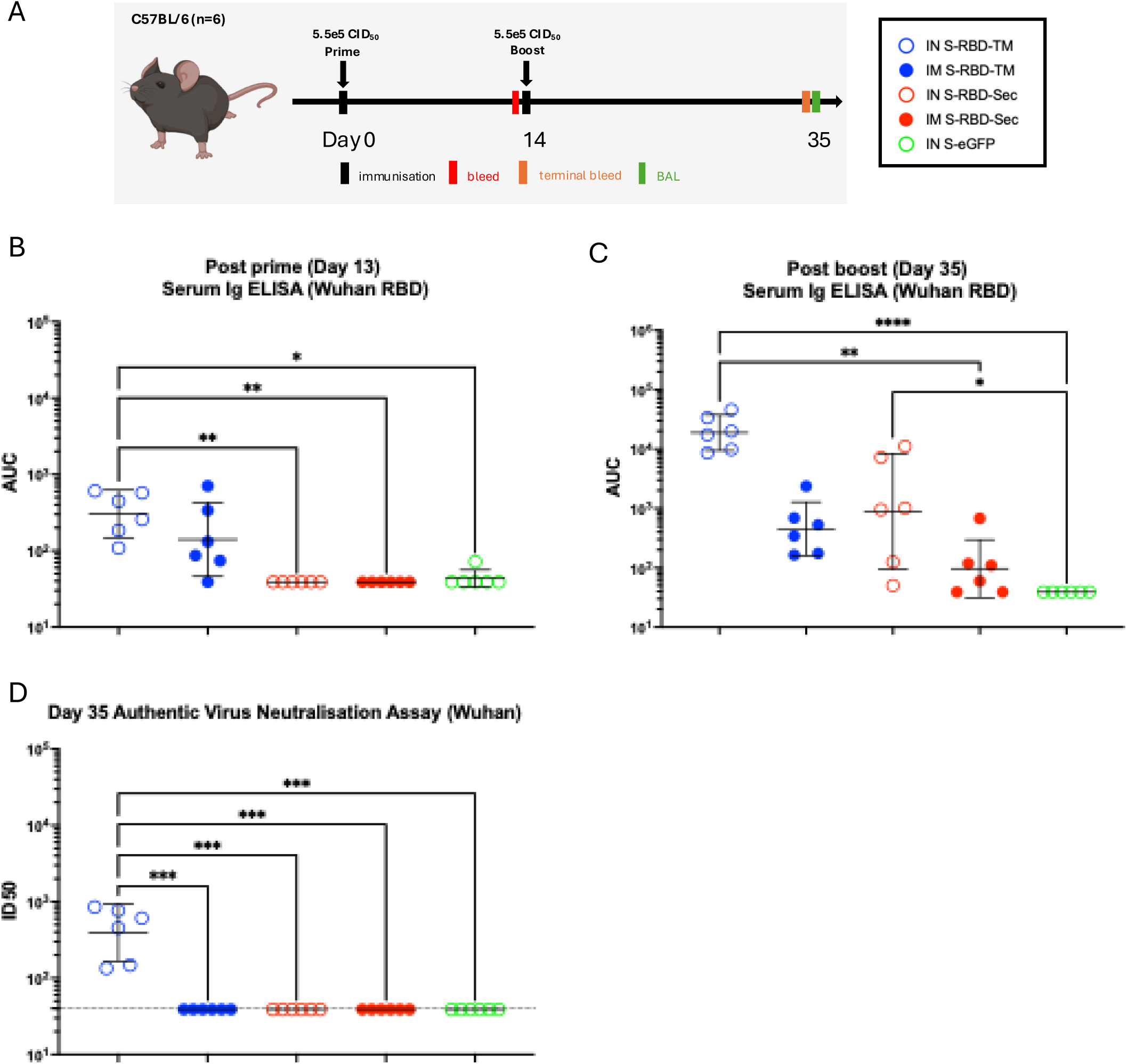
Immunogenicity of S-RBD-TM and S-RBD-Sec in mice. A. Immunisation schedule. Female C57BL/6 mice (n=6) were immunised intranasally or intramuscularly, twice, at Day 0 and Day 14 with either S-RBD-TM or S-RBD-Sec. A mock treated group of mice receiving 2 doses of S-eGFP (S-Flu expressing GFP) was included. Sera was collected on day 13 after prime and 3 weeks after boost. B. Serum antibody binding titre against protype SARS-CoV-2 Wuhan after prime. C. Serum antibody binding titre against prototype SARS-CoV-2 Wuhan after boost. D. Serum neutralising titre against authentic SARS-CoV-2 Wuhan. The values are presented as geometric mean±95% confident interval. Dotted horizontal line in D represents the lowest sample dilution tested in the assays. Values plotted as 39 in D indicating no detectable binding. Statistical difference was determined using Kruskal-Wallis test followed by Dunn’s correction test; *p<0.05, **p<0.01, ***p<0.001. (AUC: area under curve, ID_50_: half-maximal inhibitory dilution)

### Heterologous prime-boost regime of S-RBD increases breadth of antibody response against sarbecoviruses

Mounting evidence suggests that heterologous prime-boost immunisation regimes with antigenically distant antigens can induce broad immunity [56–58]. To investigate the potential of a heterologous prime-boost strategy in improving the breadth against sarbecoviruses, a second modified S-RBD-TM based on the distant Clade 3 bat sarbecovirus BM48-31 was generated using the approach described above. The titre of the virus (namely S/US/RBD-BM48-31-TM (Spike)/N1 PR8/ H3 X31) was 6.16e7 CID_50_/mL. We hypothesised that a heterologous S-RBD-TM Wuhan/ S-RBD-TM BM48-31 prime-boost regime could increase the breadth of antibody responses against mismatched sarbecoviruses across different clades and SARS-CoV-2 VOCs. Mice (female, C57BL/6) were immunised intranasally twice, 3 weeks apart, with either a homologous S-RBD-TM Wuhan prime-boost regime (homologous), a heterologous S-RBD-TM Wuhan/ S-RBD-TM BM48-31 prime-boost regime (heterologous) or with a homologous S-eGFP prime-boost regime as a control (mock) (Figure 3A). Sera and bronco-alveolar lavage (BAL) were collected 3 weeks post-boost, and the antibody response was measured. To assess whether the heterologous prime-boost regime elicited broad-spectrum antibodies against SARS-CoV-2 variants of concern (VOCs), post-boost sera were assessed by ELISA for binding to a panel of SARS-CoV-2 VOC Spike proteins (Figure 3B) and to RBDs from sarbecovirus strains (Figure 3C). The ELISA binding data are presented as line plots with significant differences between the different dosing cohorts determined by pairwise comparisons for each strain tested.

**Figure 3:**
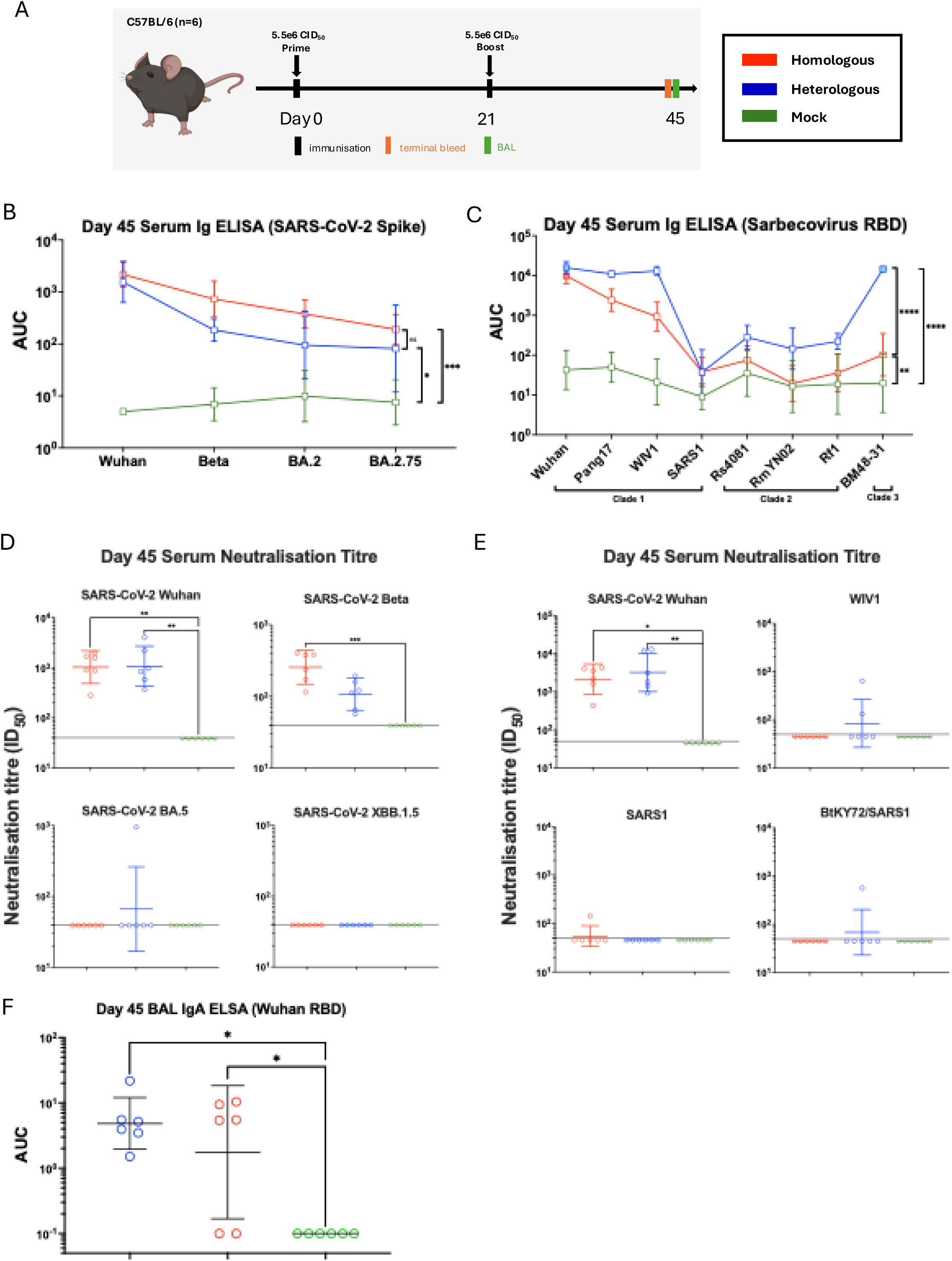
Immunogenicity of homologous and heterologous S-RBD-TM prime-boost in mice. A. Immunisation schedule. Female C57BL/6 mice (n=6) were immunised intranasally at Day 0 with S-RBD-TM Wuhan or S-eGFP control (Mock), followed by a boost at Day 21 with either S-RBD-TM Wuhan (Homologous) or S-RBD-TM BM48-31 (Heterologous). A mock treated group of mice receiving 2 doses of S-eGFP (S-Flu expressing GFP) was included. Sera was collected on day 20 after prime and 3 weeks after boost. Broncho-alveolar lavage (BAL) was collected 3 weeks after boost. B. Serum antibody binding titre against prototype SARS-CoV-2 Variants of Concern (VOCs) Spikes including Wuhan, Beta, BA.2 and BA.2.75. C. Serum antibody binding titre against sarbecovirus RBD (Clade 1 (SARS-CoV-2 Wuhan, Pang 17, WIV1, and SARS1), Clade 2: Rs4081, RmYN02 and Rf1), (Clade 3: BM48-31)). D. Serum neutralisation titre against authentic SARS-CoV-2 VOCs (Wuhan, Beta, BA.4/5, XBB.1.5). E. Serum neutralisation titre against sarbecovirus pseudovirus (Wuhan, WIV1, SARS1 and BtKY72/SARS1 hybrid). F. BAL IgA antibody binding titre against prototype SARS-CoV-2 (Wuhan). The values are presented as geometric mean±95% confident interval. Dotted horizontal lines in D and E represent the lowest serum dilution tested in the assays. Statistical difference in B and C was determined using Tukey’s multiple pairwise comparison (means of all RBD tested between immunisation cohorts) followed by the Geisser-Greenhouse correction. Values plotted at 0.1 in F indicating no detectable binding. Statistical difference in D, E and F was determined using Kruskal-Wallis test followed by Dunn’s correction test; *p<0.05, **p<0.01, ***p<0.001, ****p<0.0001. (AUC: area under curve, ID_50_: half-maximal inhibitory dilution)

Serum antibody binding to SARS-CoV-2 VOCs Spike proteins tested (Wuhan, Beta, BA.2, and BA.2.75) could be detected in both the homologous and heterologous groups, whereas no binding activity was detected in the mock treated group. No significant difference was observed between the homologous and heterologous group in terms of binding titres against all Spikes tested (Figure 3B).

We then assessed whether the heterologous prime-boost regime could increase the cross-reactivity of serum antibodies against sarbecoviruses compared to homologous prime-boost with S-RBD-TM Wuhan. RBD-binding antibody titres were measured by ELISA against a panel of 8 sarbecovirus RBDs, of which 6 were mismatched or partially matched, with BM48-31 being matched to the heterologous group. The serum binding titre to sarbecovirus RBDs of both the homologous and heterologous group were significantly higher than the S-eGFP control group (p<0.01 and p<0.0001, respectively). Notably, the heterologous group elicited a significantly higher serum binding titre to sarbecovirus RBDs compared to the homologous group (p<0.0001) (Figure 3C). The difference in serum antibody binding titre between the homologous and heterologous group was most pronounced against BM48-31, which is antigenically matched to the heterologous boost group. The heterologous group also showed increased binding to a few of the RBDs tested, such as Pang17, WIV1, and Rf1, compared to the homologous group.

We next evaluated the neutralisation potency of serum antibodies with authentic neutralisation assays against representative SARS-CoV-2 Wuhan, Beta, BA.5 and XBB.1.5 variants. No differences were seen between the homologous and heterologous groups, with both groups eliciting significantly higher neutralisation titres against SARS-CoV-2 Wuhan compared to the S-eGFP group (ID_50_: homologous: 1:1,216 and heterologous: 1:1,499, p<0.01) (Figure 3D). The homologous group elicited significantly higher serum neutralising titre against the Beta strain (p<0.001) but there was no significant difference for neutralisation against the Beta strain between the homologous and heterologous group (ID_50_: homologous:1:283, and heterologous: 1:118). No mice in the homologous group showed neutralisation against BA.5. However, 1 of the 6 mice in the heterologous group had detectable neutralising activity (ID_50_: 1:956). Neither groups showed neutralisation activity against XBB.1.5.

To assess the neutralisation potency against sarbecoviruses, *in vitro* neutralisation assays were performed using pseudoviruses (SARS-CoV-2 Wuhan, WIV1, SARS1, and BtKY72), as previously described [59] (Figure 3E). Both homologous and heterologous groups elicited significantly higher neutralisation titres against the SARS-CoV-2 Wuhan pseudovirus, compared to the S-eGFP mock control group (ID_50_: 1:2,704 and 1:5,376, p<0.05 and p<0.01, respectively). These findings are consistent with results obtained from the authentic virus neutralisation assay against Wuhan.

In contrast, cross-neutralisation against other sarbecoviruses was limited (Figure 3E). For WIV1, no neutralising activity was detected in the homologous group, while only 2 of the 6 mice in the heterologous group showed detectable neutralising activity (ID_50_: 1:644 and 1:132, respectively). Similarly, against BtKY72, all mice in the homologous group failed to elicit neutralising activity, and only one mouse in the heterologous group exhibited neutralising activity (ID_50_: 1:565). For SARS-1, a single mouse in the homologous group showed neutralisation activity (ID_50_: 1:143), whereas no mice in the heterologous group showed detectable neutralisation activity.

To assess local antibody responses in the airways, the IgA response against Wuhan RBD in BAL was measured. Both homologous and heterologous groups elicited significantly higher RBD IgA responses against the control S-eGFP group (p<0.05) (Figure 3F). The difference in IgA responses between the homologous and heterologous group was not statistically significant.

## Discussion

The emergence of SARS-CoV-2 variants and the ongoing threat of zoonotic sarbecovirus underscores the need for pan-sarbecovirus vaccines. Such vaccines could offer a proactive defence against future epidemics and pandemics. Broadly reactive vaccines may eliminate the need for strain-specific formulations, which would mark a ground-breaking shift in pandemic preparedness. The identification of numerous cross-neutralising antibodies, specifically targeting conserved RBD class 3, 4,1/4 and 5 epitopes [33, 34, 36–39, 41–43], support the feasibility of sarbecovirus RBDs as a promising target for broadly reactive vaccine strategies.

In this study, we generated and characterised two versions of S-FLU expressing the SARS-CoV-2 RBD: a soluble form (S-RBD-Sec) and a membrane-anchored form (S-RBD-TM). The use of S-FLU as a candidate vaccine offers several compelling advantages. Notably, S-FLU exhibits the capacity to infect host cells but is replication-incompetent; this controlled infection facilitates small droplet aerosol administration directly to the respiratory tract to elicit mucosal immunity [45, 60, 61]. Both S-RBD constructs were efficiently rescued using reverse genetics and demonstrated robust RBD expression in infected cells. Immunisation studies in mice revealed that both S-RBD constructs elicited systemic anti-RBD antibody responses, with S-RBD-TM showing superior immunogenicity, particularly via the intranasal route. Although T-cell responses were not evaluated in this study, we previously shown that S-FLU can elicit both local lung and systemic CD8 T-cell responses to the transgene it carries, which could further enhance the overall immunogenicity of S-RBD [45, 62].

To enhance the breadth of the antibody response, we employed a heterologous prime-boost strategy using antigenically distinct RBDs derived from the SARS-CoV-2 Wuhan strain and the Clade 3 bat sarbecovirus BM48-31. This approach broadened the binding antibody responses against sarbecoviruses, although no significant differences in neutralisation potencies were observed between homologous and heterologous regimens. This suggests that binding does not always correlate with functional neutralisation, and further optimisation may be needed to enhance cross-neutralising activity. Nevertheless, these non-neutralising antibodies could have a protective role, via Fc-mediated effector functions, as have been previously shown to aid in prevention of severe COVID-19 [63–65]. Consequently, our findings do not provide explicit evidence to suggest that the heterologous prime-boost regime has any discernible effect on breadth against mismatched SARS-CoV-2 VOCs. This is conceivable as VOCs are still closely related to the Wuhan strain (Table S1), and a heterologous boost with a distantly related virus may not enhance the breadth of responses.

There are several limitations in this study. First, we did not directly assess the protective efficacy of S-RBD-TM in a viral challenge model. Such studies could help define potential protective effects of non-neutralising antibodies. Second, tissue resident CD4+ and CD8+ T cell activity in the airways was not evaluated, which has been shown to correlate with protection and reduced disease severity against SARS-CoV-2 infection in mice [66, 67]. Lastly, although we showed robust antibody responses at 3-week post boost, the longevity of the antibody response was not determined.

In summary, our findings demonstrate the potential of S-FLU-based RBD intranasal vaccines to elicit both local and systemic antibody responses and highlight the promise of heterologous prime-boost strategies in providing pan-sarbecovirus immunity.

## Material and Methods

### Plasmids

Plasmids used for reverse genetics for S-RBD-TM production are; 6 pPol plasmids encoding the vRNA (pPol_PB1-PR8, pPol_PB2-PR8, pPol_PA-PR8, pPol_NP-PR8, pPol_NS-PR8, pPol_M-PR8, pPol_N1-PR8), 4 core initiators to supply essential viral proteins in *trans* for vRNA replication and viral packaging (pCDNA3.1_PB1-PR8, pCDNA3.1_PB2-PR8, pCDNA3.1_PA-PR8, pCDNA3.1_NP-PR8), pCDNA3.1_H1_HA-PR8 (to supply HA in trans on the producer cells’ surface) and either pPol/US/RBD (Wuhan)-TM, pPol/US/RBD (BM48-31)-TM or pPol/US/eGFP. pPol/US/eGFP had previously been synthesised for proof of principle experiments [45]. The cDNA fragment (mammalian signal sequence-BM48-31 RBD- SARS-CoV-2 Spike Transmembrane Domain flanked by *NotI* and *EcoRI* sites) was designed and synthesised by Twist Bioscience. 3,000ng of the cDNA fragment from Twist Bioscience was digested with *NotI* and *EcoRI* at 37°C for 30 min in a 40μL reaction containing 4μL of CutSmart buffer and 20 units of *NotI* and *EcoRI*, respectively. Digested cDNA fragment was purified using a QIAquick PCR purification kit. The digested cDNA fragment was then eluted in 25μL of nuclease free water, of which 6μL was incubated with 2μL of vector (PV-pPol_HA *EcoRI*-*NotI*) (previously prepared in the laboratory), 1μL T4 DNA ligase buffer and 1μL T4 DNA ligase at 25°C for 30 min. 5μL of the ligation reaction was then used to transform 10μL of *E.coli* DH5-alpha competent cells that were plated on an Ampicillin LB agar plate. The agar plate was incubated at 37°C overnight. 4 colonies were then picked and grown in LB broth overnight in a shaking incubator (200 rpm). Plasmid DNA was then extracted by alkaline lysis method using a ZymoPURE Plasmid Miniprep kit. The sequence of the extracted plasmid minipreps was confirmed using Sanger sequencing.

### Generation of S-FLU viruses

S-FLU (S-RBD) viruses were generated by reverse genetics, as described [45] with slight modifications. Briefly, 1e6 of HEK293T cells were seeded in a 6 well-plate in 2mL D10 (Dulbecco’s modified Eagle’s medium (DMEM) with 10% heat inactivated foetal bovine serum, 2mM L-glutamine, 1% penicillin and streptomycin). 24h later, media was exchanged to serum-free OptiMEM media and 1μg of each plasmid was used to transfect the HEK293T cells using the Lipofectamine2000 (ThermoFisher) transfection reagent. 4h post-transfection, media was replaced with 2mL of viral growth media (VGM; DMEM, 1% penicillin & streptomycin, 2mM L-glutamine, 0.1% BSA, 10mM HEPES) containing 1μg/mL TPCK-treated trypsin.

Approximately 72h post-transfection, the supernatant containing seed virus was harvested and clarified via centrifugation at 1400 x g for 5 min. MDCK-X31 cells (1e6/well) seeded in 6-well plate the day before were infected with the seed virus for 1h at 37°C. Media was then replaced with 2mL VGM containing 1μg/mL TPCK-treated trypsin 1h post-infection and plates were incubated at 37°C. The propagated virus (S-RBD-TM) was harvested 48h post-infection. The S-RBD-TM was further propagated by infecting the 2mL (diluted in 10mL of VGM) of virus in a T175 seeded with 5e6 MDCK-X31 the day before. After 1h of incubation at 37°C, media was replaced with 40mL of fresh VGM containing 1ug/mL of TPCK-treated trypsin. After 48h, supernatant containing the virus was harvested and clarified via centrifugation at 1400 x g for 5 min to pellet any cellular debris. Harvested virus was aliquoted and stored at −80°C.

### Virus titration (CID_50_)

Viral titres were determined by fifty percent cell culture infectivity dose (CID_50_) in MDCK-SIAT1 indicator cells as previously described with slight modifications. Briefly, MDCK-SIAT1 cells (3×10^4^/well) seeded in a flat-bottomed 96-well plate were infected with harvested virus in a 2-fold dilution series. After overnight incubation at 37°C, the media was removed, and 100µL of 10% formalin (4% paraformaldehyde) in PBS was added at 4°C for 30 min. The plates were washed twice with PBS and stained with 50μL of 10μg/mL of EY-6A (anti-RBD human IgG1 monoclonal antibody (binds to all SARS-like betacoronavirus RBD)[68] in PBS/0.1% BSA and incubated for 1h at RT on a plate shaker. For S/US/RBD-Sec/N1 PR8/H3 X31, infected cells were permeabilised for 20min at RT with PBS/ 0.5% Triton-X/ 20mM glycine prior to staining. The primary antibody was then removed, and the wells were washed twice with PBS and incubated for 1h on a plate shaker with 50μL of Goat-Anti-Human (GAH) AF647 (1:500) in PBS/0.1% BSA. After washing with PBS, 100µL of 1% formalin (0.4% paraformaldehyde) was added to each well, and plates were read using a Clariostar spectrophotometer (BMG Labtech). The CID_50_/mL was calculated as previously described [51, 52], and the titres are: S/US/RBD-Sec/N1 PR8/H3 X31: 7.68e7 CID_50_/mL, S/US/RBD-TM/N1 PR8/H3 X31:1e7 CID_50_/mL, S/US/RBD-BM48-31-TM (Spike)/N1 PR8/ H3 X31: 6.16e7 CID_50_/mL.

### RBD expression confirmation *in vitro*

MDCK-SIAT1 cells (1e6/well) were cultured overnight in 6-well plates with 2mL of D10 at 37°C. Media was then replaced with 0.5e6 CID_50_ of virus in 1mL of VGM and incubated overnight at 37°C. After removal of the media, 1mL of 10% formalin was added and incubated at 4°C for 30 min. The plate was washed twice with 1mL PBS, then EY-6A IgG (10 µg/mL in 1mL PBS/0.1% BSA) was applied and incubated at RT for 1h. After washing, Goat Anti-Human-AF647 (1:500 in 1mL PBS/0.1% BSA) was added, and the plate was incubated for 1h at RT. Following two washes with 1mL PBS, 1mL of 1% formalin was added per well for fluorescence microscopy.

### Mice and immunisation

Mice used in this study were purchased from Envigo Ltd., UK and procedures carried out at the Biomedical Services Unit (BMS) (University of Oxford, United Kingdom) in accordance with the UK Animals (Scientific Procedures) Act 1986 and with approval from the local Animal Welfare and Ethical Review Body (AWERB) (Project Licence PP9362617). Mice were housed in accordance with the UK Home Office ethical and welfare guidelines and fed on standard chow and water *ad libitum*. Female C57BL/6 mice at 4-6 weeks of age were used in this study. For intranasal delivery, mice were lightly anaesthetised with inhaled isoflurane and 50μL of S-RBD was delivered via the nostrils. For intramuscular delivery, mice were anaesthetised with inhaled isoflurane and 50μL of S-RBD was injected using an insulin needle into the thigh muscle. Post prime blood was collected via the tail vein using a capillary tube. Mice were euthanized by gradual chamber fill with compressed CO₂ (displacement rate: 30–70% of chamber volume per minute). Sera were collected by cardiac puncture, with whole blood drawn into Microtainer SST tubes (BD). Samples were clotted at room temperature for 1h, centrifuged at 10,000 × g for 5 min, and the clarified sera were heat-inactivated at 56 °C for 30 min before storage at −20 °C. To collect BAL, the trachea of euthanised mice was exposed by blunt dissection, and a small incision was made to insert a cannula (0.75 mm, Harvard Apparatus, UK) attached to a 21G needle. The cannula was advanced until positioned just above the carina. A 1 mL syringe was used to instil PBS into the lungs, followed by gentle aspiration to recover the fluid. This procedure was repeated twice, and the collected lavage was stored at −20 °C until further processing.

### Production of soluble Spike and RBD for ELISA

The SARS-CoV-2 and VOCs Spikes (HexaPro) (Wuhan, Beta, BA.2 and BA.2.75) were previously produced in the laboratory using the method described below. The plasmids encoding soluble sarbecovirus RBD (RBD-6H or RBD-ST1 (Wuhan), BM48-31-AviTag, Pang17-AviTag, WIV1-AviTag, Rs4081-AviTag, RmYN02-AviTag, Rf1-AviTag) were expressed in Expi293 cells via transient transfection according to the manufacturer’s instructions. At day 6-7 post-transfection, expressed proteins were purified from culture supernatants via immobilised nickel affinity chromatography (IMAC) using an automated protocol implemented on an AKTAStart system (GE Healthcare). Affinity-purified proteins were diafiltered and concentrated into PBS using 10k MWCO Amicon spin filters. Protein concentrations were determined from A_280_ measurements using a Nanodrop Spectrophotometer (ThermoFisher).

### Serum IgG ELISA

ELISAs were conducted as previously described [53]. Briefly, 50μL of 2μg/mL of purified RBD or 1μg/mL of trimeric Spike (HexaPro)[69] proteins diluted in PBS were added to each well of Maxisorp NUNC plates (ThermoFisher) overnight at 4°C or at RT for 1-2h on a plate shaker. Plates were washed 3 times with PBS and blocked with 300μL of 5% skim milk in PBS for 1h at RT on a plate shaker or overnight at 4°C. In round bottomed 96-well plates, mouse sera were diluted (starting dilution 1:100) in PBS/0.1% BSA in a half-log serial dilution in duplicate. NUNC plates were washed with PBS and 50μL of the diluted sera was transferred to the coated NUNC plates for 1h at RT before being washed with PBS 3 times. The secondary HRP conjugated goat anti-mouse immunoglobulin antibody (Dako P0447) was diluted 1:800 in PBS/0.1% BSA, and 50μL were added into each well of the NUNC plate for 1h at RT. Plates were then washed and developed with 50μL of KPL SureBlue substrate for 5 min and stopped with 50μL of 1M H_2_SO_4_. Absorbance (OD_450_) was measured on a Clariostar spectrophotometer (BMG Labtech). ELISA titres were calculated and presented as Area Under Curve (AUC).

### Authentic virus microneutralisation assays (SARS-CoV-2 VOCs)

Microneutralisation assays against authentic SARS-CoV-2 viruses were conducted as described [54, 55]. Triplicate serial dilutions of sera (20μL) were pre-incubated with 100 focus-forming units (FFUs) of virus for 30 min at RT. After pre-incubation, 100μL Vero CCL81 cells (4.5×10^4^) were added and incubated at 37°C, 5% CO_2_. 2h later, 100μL of a 1.5% carboxymethyl cellulose-containing overlay was added to prevent satellite focus formation. 20h (Wuhan), 18h (Beta) and 22h (Omicron BA.5 and XBB.1.5) post-infection, monolayers were fixed with 4% paraformaldehyde at RT for 30 min, permeabilised with 100μL 2% Triton X-100 for 30 min at 37°C and stained for the nucleoprotein with FB-9B. After development with a peroxidase-conjugated antibody and TrueBlue peroxidase substrate, infectious foci were enumerated by ELISpot reader. Half-maximal inhibitory dilutions (ID_50_) were assessed using 4-parameter nonlinear regression in GraphPad Prism 10. Assays were performed in a containment level three facility under a license from the Health and Safety Authority, UK.

### Pseudovirus neutralisation assays (sarbecoviruses)

Pseudoviruses based on a lentiviral vector were prepared as described [35, 70] using genes encoding S protein sequences lacking C-terminal residues in the cytoplasmic tail: 21 residue (SARS-CoV-2 Wuhan and WIV1) or 19 residue cytoplasmic tail deletions (SARS-CoV, BtKY72/SARS-1 chimera). For neutralisation assays, pseudovirus was incubated with three-fold serially diluted sera from immunised mice for 1h at 37°C, then the serum/virus mixture was added to 293T_ACE2_ target cells. After 48h incubation, media was removed, and cells were lysed with Britelite Plus reagent (Perkin Elmer), and luciferase activity was measured as relative luminesce units (RLUs). Relative RLUs were normalised to RLUs from cells infected with pseudovirus in the absence of serum. Half-maximal inhibitory dilutions (ID_50_) were derived using 4-parameter nonlinear regression in AntibodyDatabase [71].

### Statistical Analysis

Pairwise comparisons were performed (Figure 3B and 3C), as described previously [72], to evaluate sets of binding titre against individual RBD across different immunisation cohorts and to determine whether results between cohorts were significantly different. Statistical differences in ELISA and neutralisation titres between immunised groups were assessed using analysis of variance (Kruskal-Wallis), followed by followed by Dunn’s correction test, pairing by Spike/RBD protein or viral strain. Statistical analysis was performed using GraphPad Prism version 10. Statistical significance is represented as *p<0.5, **p<0.01, ***p<0.001, **** p<0.0001.

## Author Contributions

Conceptualisation: T.K.T, A.R.T. Methodology: J.M., T.K.T. Investigation: J.M., T.K.T., D.M., L.S., M.A., A.C., J.R.K. Supervision: A.R.T., T.K.T. Writing (original draft): J.M., T.K.T. Writing (review and editing): T.K.T, W.S.J., J.R.K, P.J.B., A.R.T.

## Funding

The work was funded by the Townsend-Jeantet Charitable Trust (charity number 1011770), the COVID Response Fund (University of Oxford), and the National Institutes of Health (P01-AI165075 to P.J.B.).

## Acknowledgments

We would like to express our gratitude to the staff members at the Biomedical Science Services, John Radcliffe Hospital (Oxford University) for their assistance with the mouse experiments. This manuscript is the result of funding in whole or in part by the National Institutes of Health (NIH). It is subject to the NIH Public Access Policy. Through acceptance of this federal funding, NIH has been given a right to make this manuscript publicly available in PubMed Central upon the Official Date of Publication, as defined by NIH.

---

**Figure S1:**
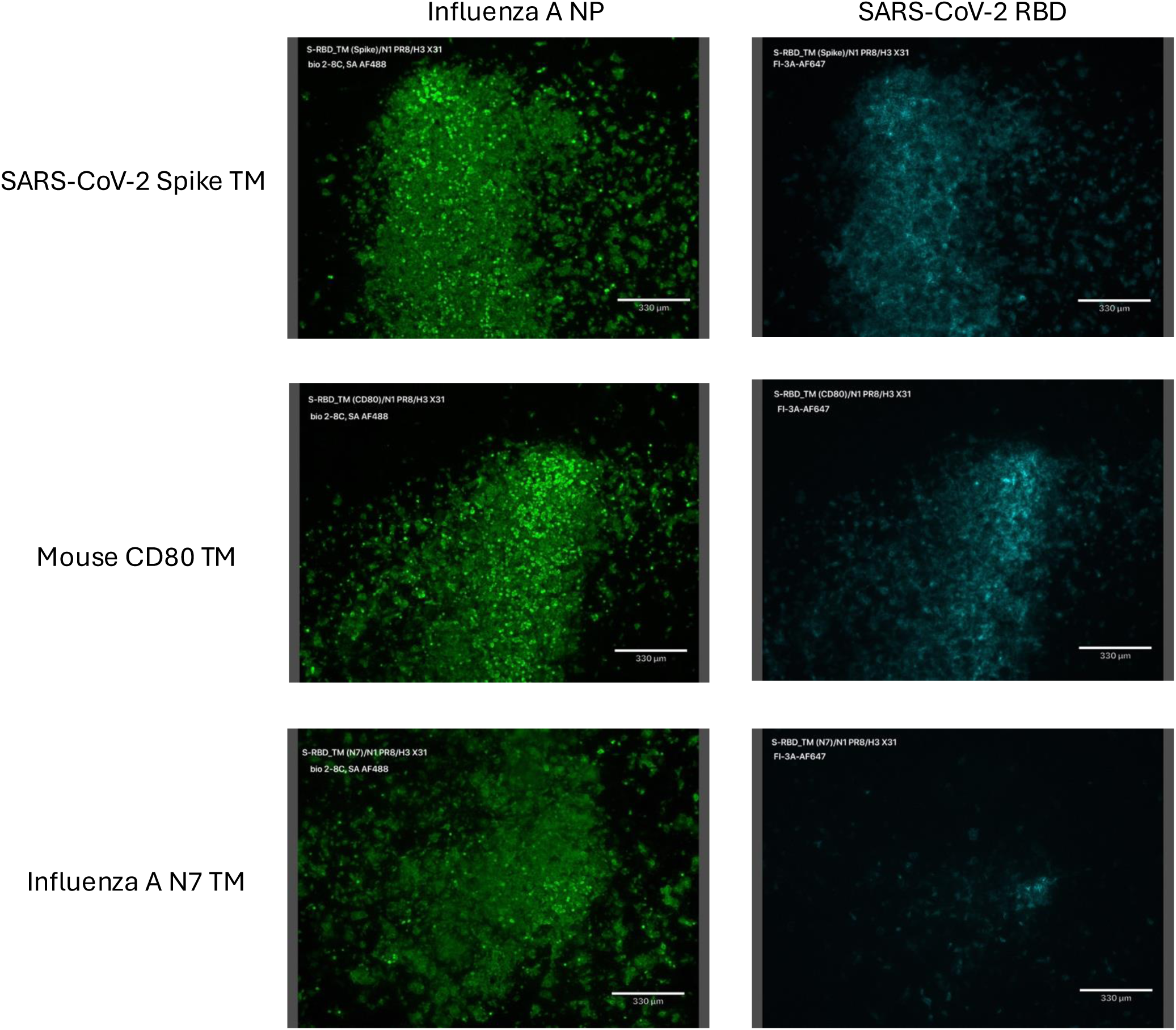
Expression of influenza NP and RBD in cell culture infected with S-RBD-TM with various TM. MDCK cells seeded in 6-well plate were infected with S-RBD-TM (fused to a SARS-CoV-2 Spike TM, mouse CD80 TM or an influenza neuraminidase N7 transmembrane domain). 16 hours post infection, cells were formalin fixed, permeabilised with 0.5% Triton-X 100 and stained with an RBD-specific IgG1 (FI-3A) labelled with AF647, and a biotinylated anti-Influenza NP IgG1 (2-8C biotin), followed by Streptavidin labelled with AF488. Stained cells were visualised using a fluorescence microscope. Scale bar: 330uM. (TM: transmembrane domain, AF647: Alexa Fluor 647, AF488: Alexa Fluor 488)

**Table S1:** Position-by-position mutation matrix of the SARS-CoV-2 spike RBD (residues 319–541). The table shows the specific amino acid changes in major variants relative to Wuhan SARS-CoV-2. Each column represents an RBD residue and each row a variant. Cells list the observed amino acid substitution at each position; blank cells indicate identity with the reference sequence.

| SARS-CoV-2 Variant | G339 | R346 | S371 | S373 | S375 | K417 | N440 | G446 | L452 | N460 | S477 | T478 | E484 | F486 | Q493 | G496 | Q498 | N501 | Y505 |
| --- | --- | --- | --- | --- | --- | --- | --- | --- | --- | --- | --- | --- | --- | --- | --- | --- | --- | --- | --- |
| Wuhan | - | - | - | - | - | - | - | - | - | - | - | - | - | - | - | - | - | - | - |
| Alpha (B.1.1.7) | - | - | - | - | - | - | - | - | - | - | - | - | - | - | - | - | - | N501Y | - |
| Beta (B.1.351) | - | - | - | - | - | K417N | - | - | - | - | - | - | E484K | - | - | - | - | N501Y | - |
| Gamma (P.1) | - | - | - | - | - | K417T | - | - | - | - | - | - | E484K | - | - | - | - | N501Y | - |
| Delta (B.1.617.2) | - | - | - | - | - | - | - | - | L452R | - | - | T478K | - | - | - | - | - | - | - |
| Omicron BA.1 | G339D | - | S371L | S373P | S375F | K417N | N440K | G446S | - | - | S477N | T478K | E484A | - | Q493R | G496S | Q498R | N501Y | Y505H |
| Omicron BA.2 | G339D | - | - | - | - | K417N | N440K | - | - | - | S477N | T478K | E484A | - | Q493R | - | Q498R | N501Y | - |
| Omicron BA.4 / BA.5 | G339D | - | - | - | - | K417N | N440K | - | L452R | - | S477N | T478K | E484A | F486V | Q493R | - | Q498R | N501Y | - |
| BA.2.75 | G339D | - | - | - | - | - | N440K | G446S | - | - | S477N | T478K | E484A | - | Q493R | - | Q498R | N501Y | - |
| XB B / XB B.1.5 | G339D | R346T | - | - | - | - | - | G446S | - | N460K | - | - | - | - | - | - | - | - | F486P / F486S |

## Notes

### Competing Interest Statement

The authors have declared no competing interest.

